# rePROBE: Workflow for Revised Probe Assignment and Updated Probe-Set Annotation in Microarrays

**DOI:** 10.1101/2020.03.10.985119

**Authors:** Frieder Hadlich, Henry Reyer, Michael Oster, Nares Trakooljul, Eduard Muráni, Siriluck Ponsuksili, Klaus Wimmers

## Abstract

Commercial and customized microarrays are valuable tools for the analysis of holistic expression patterns, but require the integration of the latest genomic information. This study provides a comprehensive workflow implemented in an R package (rePROBE) to assign the entire probes and to annotate the probe sets based on up-to-date genomic and transcriptomic information. The rePROBE R package is freely available at https://github.com/friederhadlich/rePROBE. It can be applied to available gene expression microarray platforms and addresses both public and custom databases. The revised probe assignment and updated probe-set annotation were applied to commercial microarrays available for different livestock species, *i.e.* ChiGene-1_0-st (*Gallus gallus, 443,579 probes*; *18,530 probe sets*), PorGene-1_1-st (*Sus scrofa, 592,005*; *25,779*) and BovGene-1_0-st (*Bos taurus, 530,717*; *24,759*) as well as human (*Homo sapiens*, HuGene-1_0-st) and mouse (*Mus musculus*, HT_MG-430_PM) microarrays. Using current specie-specific transcriptomic information (RefSeq, Ensembl and partially non-redundant nucleotide sequences) and genomic information, the applied workflow revealed 297,574 probes for chickens (pig: 384,715; cattle: 363,077; human: 481,168; mouse: 324,942) assigned to 15,689 probe sets (pig: 21,673; cattle: 21,238; human: 23,495; mouse: 32,494). These are representative of 12,641 unique genes that were both annotated and positioned (pig: 15,758; cattle: 18,046; human: 20,167; mouse: 16,335). Additionally, the workflow collects information on the number of single nucleotide polymorphisms (SNPs) within respective targeted genomic regions and thus provides a detailed basis for comprehensive analyses such as quantitative trait locus (eQTL) expression studies to identify quantitative and functional traits.

## Introduction

Current breeding goals and selection criteria for livestock species go beyond performance and carcass parameters and also consider the variation of functional traits [1,2]. To identify genetically robust farm animals, respective experimental approaches often require information from the various ‘omics’ levels [3]. In fact, the use of expression data enables the generation or the verification of hypotheses on biological processes. Microarray experiments are a valuable tool for obtaining precise information about co-expressed genes and gene networks, interactions between genes and traits, and to understand phenotypic variation. In fact, the large amount of publicly available biological data from microarray analyses stored in the Gene Expression Omnibus (GEO) [4] can also be used to generate new hypotheses.

Microarrays are essentially a collection of single-stranded DNA oligos called ‘probes’. For Affymetrix arrays, each probe counts 25 bases, which should be complementary to predefined genomic target regions. Approximately 15-27 probes are aggregated to ‘probe sets’ to represent a certain transcript (main probes). Moreover, up to 30,000 control probes are used to verify amplification and hybridization (spiked-in bacteria probes) making the microarray a reliable tool for holistic transcriptomic analysis. However, the need to clarify the identity of microarray probes and probe sets has been demanded elsewhere [5,6]. Holistic expression analyses require highly accurate annotation data to ensure the quality and reliability of a data set. In fact, microarray annotations are used to be updated routinely [7−9], since there is a steadily growing body of genomic knowledge including coding and non-coding sequences, genomic variations, gene functions, gene localizations, and regulatory mechanisms.

Nowadays, the RNA-sequencing approach employing next-generation-sequencing technologies has become popular for genome-wide transcriptome profiling [10]. Being an open system that offers opportunities to discover novel splice variants and certain fractions of the RNA [11], the microarray platform is a closed system based on existing knowledge about the genome sequence. As such, microarrays represent a high-throughput and labour-saving approach in combination with a robust out-of-the-box data analysis. For processing and downstream analyses, powerful tools have been developed to facilitate identification of differentially expressed genes and insights into functional enrichment [12,13]. The combination of RNA-sequencing and microarray approaches is thus valued to obtain a comprehensive picture at the expression level [14].

Keeping pace with the current development in genomic research, we aimed to provide a workflow enabling (i) a re-assignment of microarray probes including the correction for existing SNPs and (ii) a user-friendly update based on the current reference annotation data.

Specifically, any combination of public and custom databases with different ranking priorities was implemented. The developed workflow is based on a publicly available R [15] package termed rePROBE and is described in detail for transcriptome analysis on the example of commercial microarrays of chicken, pig, cattle, human and mouse, but is not limited to these.

## Methods

In the first step of the workflow, the analysis focused on the exclusion of a number of probes from the initial probe to probe-set assignment due to (i) multiple matches to chromosomal regions or (ii) mismatches to genomic sequences. In the second step, the gene symbol annotation of probe sets was updated. To demonstrate the feasibility of the workflow, commercially available microarrays were reassigned and re-annotated. Thus, ChiGene-1_0-st (*Gallus gallus*; 443,579 probes; 18,530 probe sets), PorGene-1_1-st (*Sus scrofa*; 592,005 probes; 25,779 probe sets), BovGene-1_0-st (*Bos Taurus*; 530,717 probes; 24,759 probe sets), HuGene-1_0-st (*Homo sapiens*; 824,740 probes; 28,869 probe sets) and HT_MG-430_PM (*Mus musculus*; 496,468 probes; 45,101 probe sets; all Affymetrix, Santa Clara, CA) microarrays were used. Here, respective amplification and hybridization controls were omitted. The exemplary procedure employed four different databases, two of which have a global nature (RefSeq and Ensembl) and the other two represent additional data sources (Nucleotide collection [NT] and RefSeq DNA database [DNA]). Regarding the assignment of probes to probe sets, the workflow allows a ranking of the database used to define the priority level for each database. Accordingly, the same ranks are processed simultaneously, whereby a successful mapping excludes probes for subsequent rank processing. For chicken, pig and cattle examples, RefSeq and Ensembl were given rank 1 while NT and DNA were assigned rank 2. For human and mouse arrays solely DNA was assigned rank 2.

The workflow offers the opportunity to implement one or more public and custom databases in a user-friendly manner. For the species of interest, specific files for annotation, single nucleotide polymorphism (SNP) variation and genome reference were automatically retrieved using the rePROBE *prepare_data* function. Corresponding databases include RefSeq (Gallus_gallus-5.0 Release 103, Sscrofa11.1 Release 106, Bos_taurus_UMD_3.1.1 Release 105, GRCh38.p12 Release 108, GRCm38.p4 Release 107; http://ftp.ncbi.nih.gov/genomes) and Ensembl (Gallus_gallus.GRCg6a, Sus_scrofa.Sscrofa11.1, Bos_taurus.ARS-UCD1.2, Homo_sapiens.GRCh38.p12, Mus_musculus.GRCm38.p6; all Release 97; ftp://ftp.ensembl.org/pub). Alternatively, FASTA sequences obtained from the Nucleotide collection (NT; accessed for chicken, pig and cattle on 12 July 2019, http://www.ncbi.nlm.nih.gov/ncbisearch/; sequence type: nucleotide) and Ensembl DNA database including corresponding SNP information from dbSNP were obtained and processed. This feature is also implemented in the *prepare_data* function. Data retrieved from NT were restricted to the respective organism and included sequences ranging between 50 and 10,000 nucleotides.

The applied workflow for the probe classification using four different databases and two ranks is schematically shown in **Figure 1**. All mapping analyses were performed with the R package Rbowtie (v1.24.0) using the Bowtie short read aligner as originally presented by Langmead *et al.* [16]. Parameters were set to ‘-y --best --strata –a’ and ‘v=2’, allowing up to two mismatches. Firstly, initial probe sequences were mapped to sequences retrieved from rank 1 database(s). If multiple databases are used, the individual genomic reference can be defined (*e.g.* RefSeq, Ensembl). Mapping results that exhibited the identical genomic position (e.g. transcript variants) were aggregated. Furthermore, probes assigned to different genomic positions were discarded. After correcting for SNPs, perfect matching probes were considered unique and specific in terms of rank 1.

**Figure 1.**
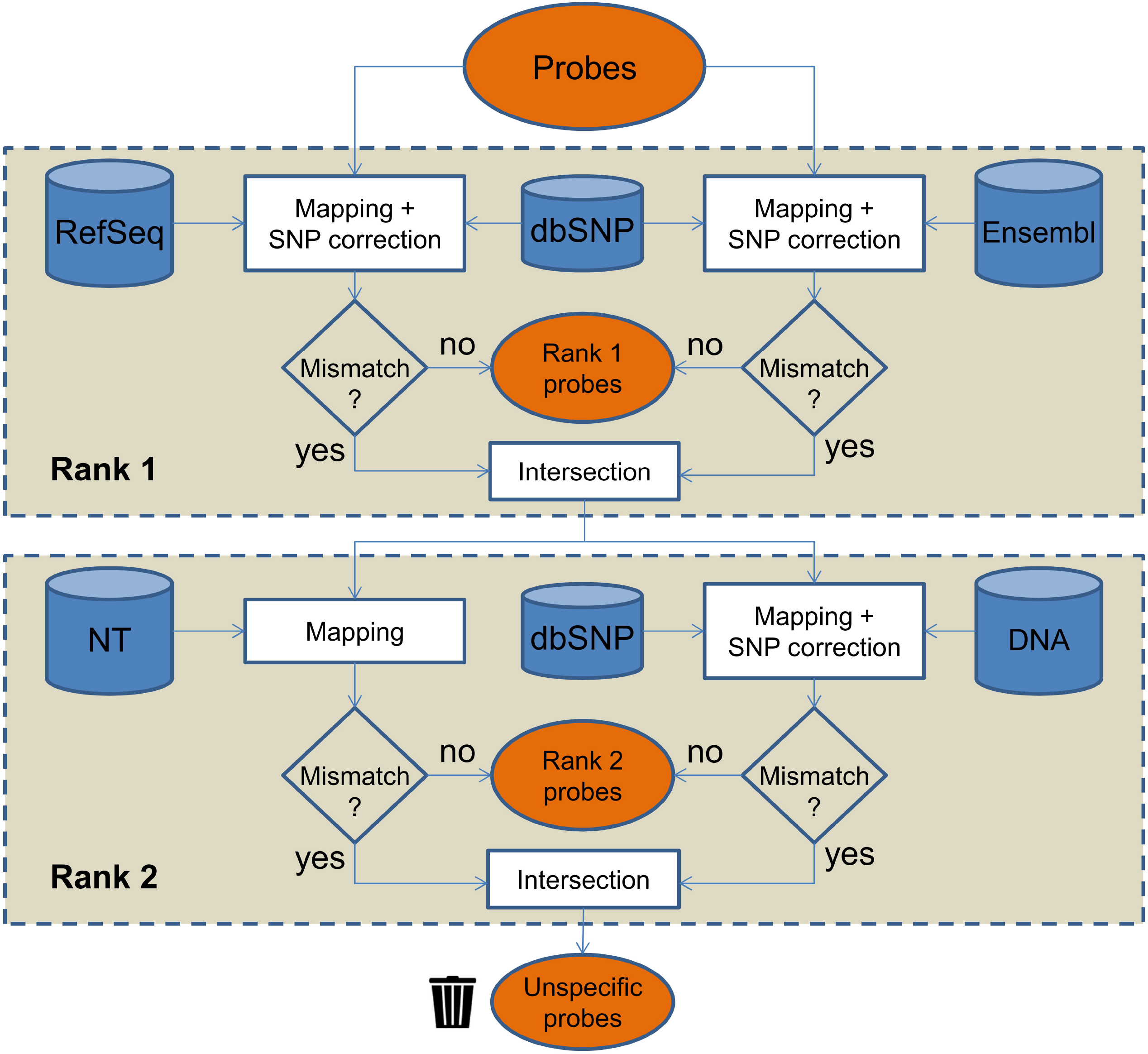
Workflow to analyse probe sequences combining information from four different databases. All probes were initially mapped against RNA sequences retrieved from RefSeq and Ensembl. Mapping results were corrected for known SNPs using information retrieved from dbSNP. Probes without mismatch in any tested RNA database were considered *transcript-specific* probes. Remaining probes which perfectly mapped to the DNA sequences or any NT database-derived sequence were considered *genome-specific* probes. Probes which failed to be mapped to any of the source sequences were discarded and considered *unspecific* probes.

The information obtained from the *prepare_data* function was used to revise the assignment of probes to probe sets and, ultimately, to update the annotation of probe sets by using the rePROBE *run* function. Specifically, generated mapping information retrieved for rank 1 was evaluated as depicted in **Figure 2** (left panel). If the probes of rank 1 represent ≥ 50% of the probes initially assigned in the respective probe set, the remaining probes were discarded and will not be processed in terms of rank 2 databases. Probes which have not been successfully assigned according to rank 1 processing were mapped to NT sequences allowing no mismatches (‘v=0’ due to missing SNP information) and to DNA sequences (Figure 1, lower panel) allowing SNP correction of up to two mismatches. Reverse orientation of sequences was allowed only for DNA entries. Perfect alignments were considered rank 2 probes. Probes which failed to be aligned to any of the source sequences or which are defined as uninformative based on the probe-to-probe-set assignment considered unspecific probes. Hence, an initially defined probe set might (i) contain entirely rank 1 specific probes, (ii) contain rank 2 information to approximate the best-choice assignment, or (iii) be completely removed. For the annotation of probes and probe sets, current information was retrieved from the corresponding databases and compiled in comprehensive probe and probe-set annotation files, suitable for numerous subsequent analyses.

**Figure 2.**
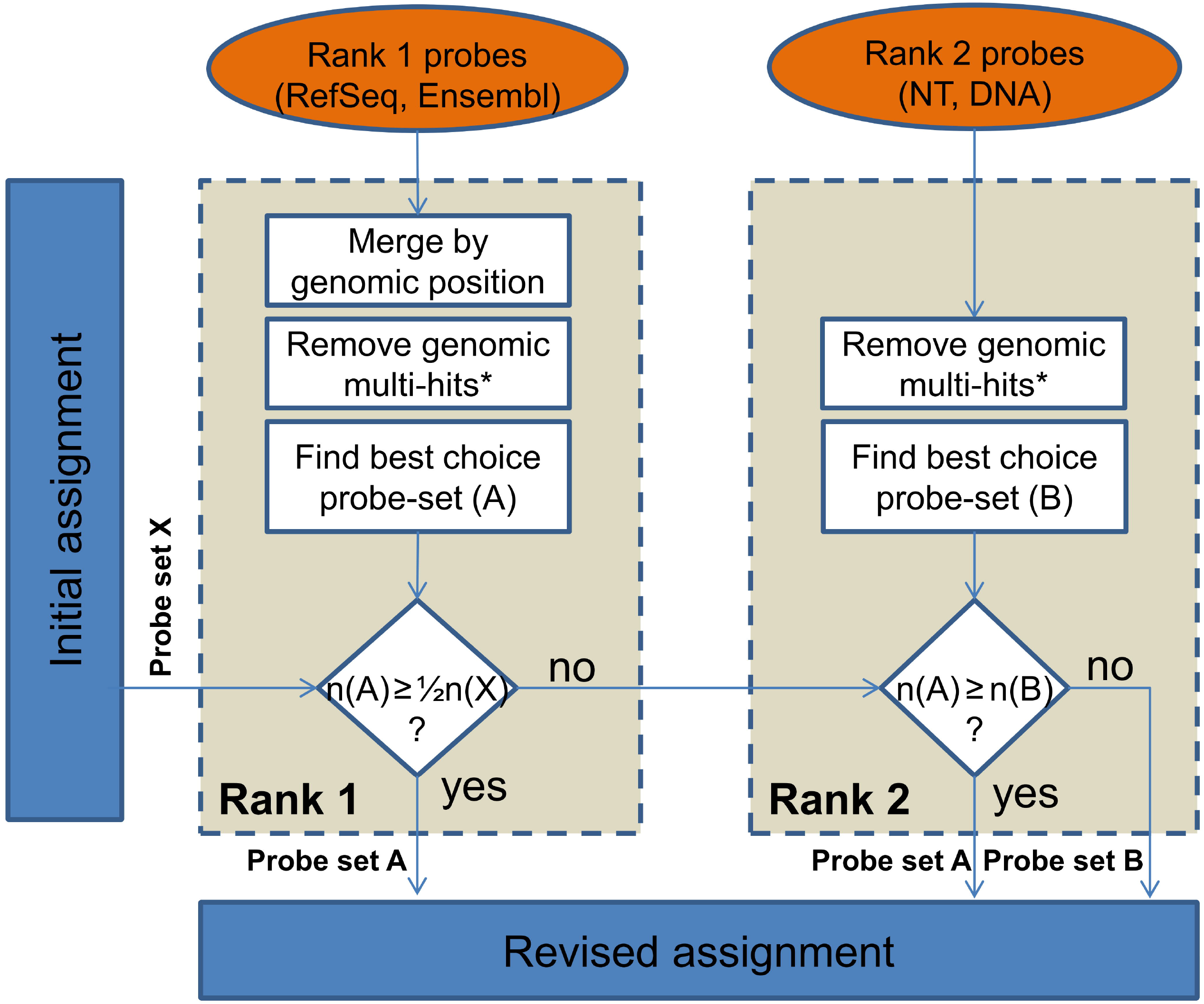
Workflow applied to each probe set. Each of the probe sets obtained from the initial assignment was processed individually (probe set X). The first step comprised only transcript-specific probes of each probe set which were uniquely mapped to the exonic part of the genome. The transcript variants were merged by genomic position. Sharing a single gene annotation prompted an assignment as probe set A (prominent role of gene symbols). Transcript-specific probes were used in the revised assignment if at least 50 % of the probes initially assigned to a certain probe set were present. Otherwise, probes with only unique genomic mappings to any alternative database were assigned as probe set B containing either a NT annotation or, if more dominating, a unique DNA region. Subsequently, the dominating probe set (containing more probes, *i.e.* either probe set A or probe set B) was used for the revised assignment. *genomic multi-hit: only applied for known chromosomes (1, 2, …, MT, X, Y).

## Application and reporting formats

The assignment of successfully mapped probes is shown in **Figure 3**. Specifically, chicken mapping information retrieved from various databases revealed 89.8% of the spotted probes to be assigned as rank 1 probes (pig: 89.4%; cattle: 92.2%; human: 93.2%; mouse: 76.3%); 87.2% of these probes matched transcript sequences retrieved from both RefSeq and Ensembl (pig: 88.9%; cattle: 90.0%; human: 86.3%; mouse: 85.7%). Ultimately, the revised probe assignment which keeps only probes of best choice probe sets relies on 67.1% of all given probes for the chicken chip (pig: 65.0%; cattle: 68.4%; human: 58.3%; mouse: 65.5%). The revised assignment therefore used a subset of the spotted information on the various microarrays. The reliability of the information obtained at the probe level has therefore been considerably improved. This probe information includes details of the genomic mapping position, exon position, and the number of putative SNP positions. The integration of these data at the probe level enables the usage of the microarrays for in-depth analyses such as the investigation of transcript and splice variants.

**Figure 3.**
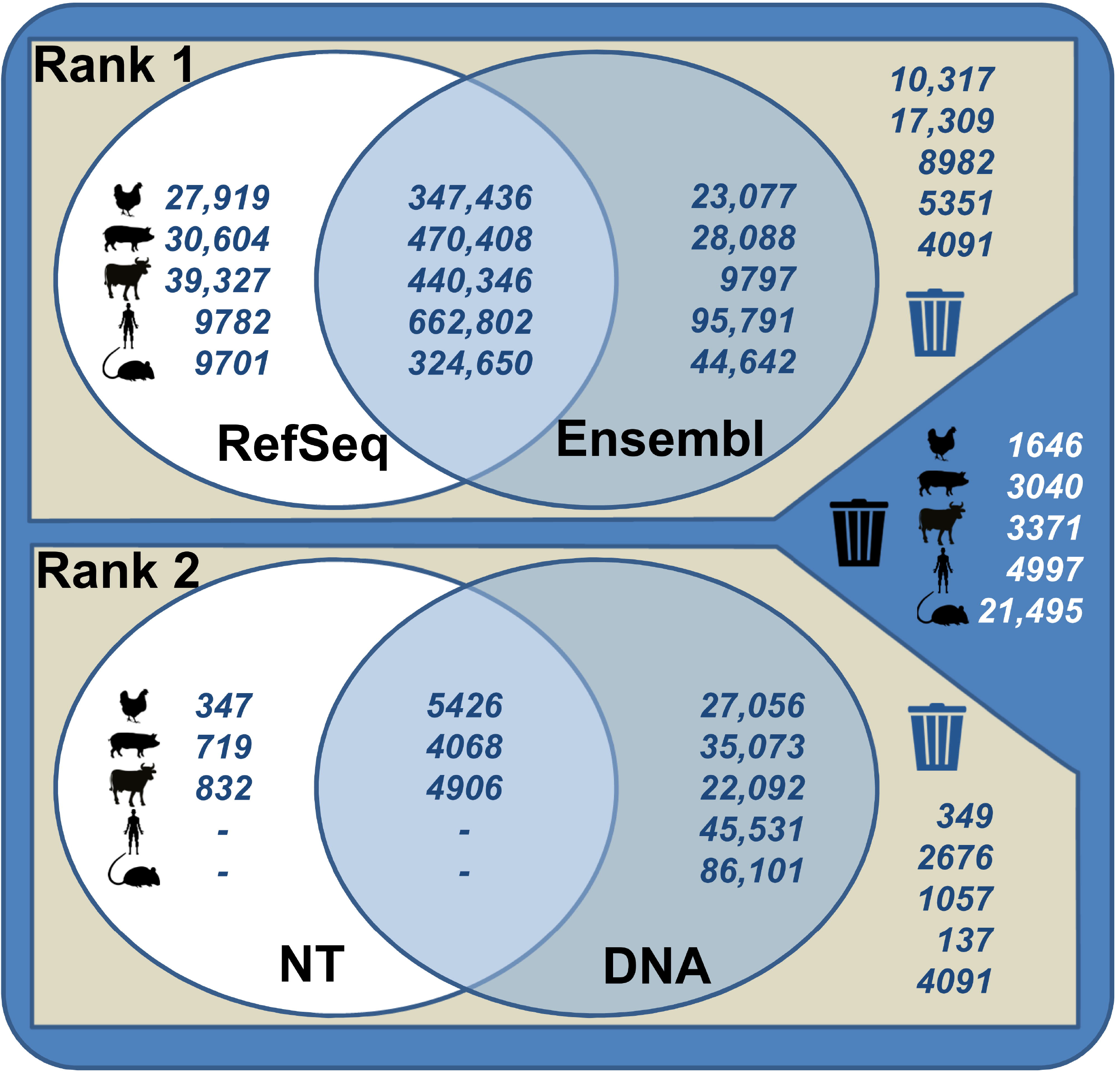
Probe mapping numbers using specific genomic databases for chickens, pigs, cattle, human and mouse microarrays. Generated mapping information retrieved for rank 1 (RefSeq, Ensembl) and rank 2 databases (Nucleotide collection [NT], DNA) were displayed as Venn diagrams. For the exemplary microarrays, the mapping revealed a number of probes which were identified by one or two databases as well as probes which failed to be aligned to any of the source sequences.

The revised probe-set assignment substantially benefits from the available genomic information to ensure the measurement of specific targets. In the chicken microarray, 12,581 probe sets remained containing ≥50% of their initially affiliated probes (pig: 17,374; cattle: 17,199; human: 18,036; mouse: 30,050). A total of 8815 probe sets were entirely unaffected and correspond to the initial assignment (pig: 10,661; cattle: 13,074; human: 13,335; mouse: 23,522). However, the revised probe to probe-set assignment identified 2841 probe sets which comprised exclusively unspecific probes (pig: 4106; cattle: 3521; human: 5601; mouse: 12,607).

According to current genomic knowledge, the applied workflow for the chicken microarray revealed 15,689 probe sets representing 12,641 unique genes (pig: 21,673; 15,758; cattle: 21,238; 18,046; human: 23,495; 20,167; mouse: 32,494; 16,335) to be both annotated and positioned. The respective gene symbols were retrieved from (i) both RefSeq and Ensembl (chicken: 8159; pig: 11,675; cattle: 12,563; human: 11,540; mouse: 18,497), (ii) RefSeq (chicken: 3306; pig: 3948; cattle: 5157; human: 3319; mouse: 912), (iii) Ensembl (chicken: 1870; pig: 2662; cattle: 1129; human: 6123; mouse: 4536), (iv) both NT and DNA (chicken: 478; pig: 426; cattle: 395), and (v) NT (chicken: 36; pig: 84; cattle: 58) entries. The number of probe sets from which both genomic position (**Figure 4A**) and annotation (**Figure 4B**) are currently known reflects the improvement in accessible genomic knowledge since the initial microarray design.

**Figure 4.**
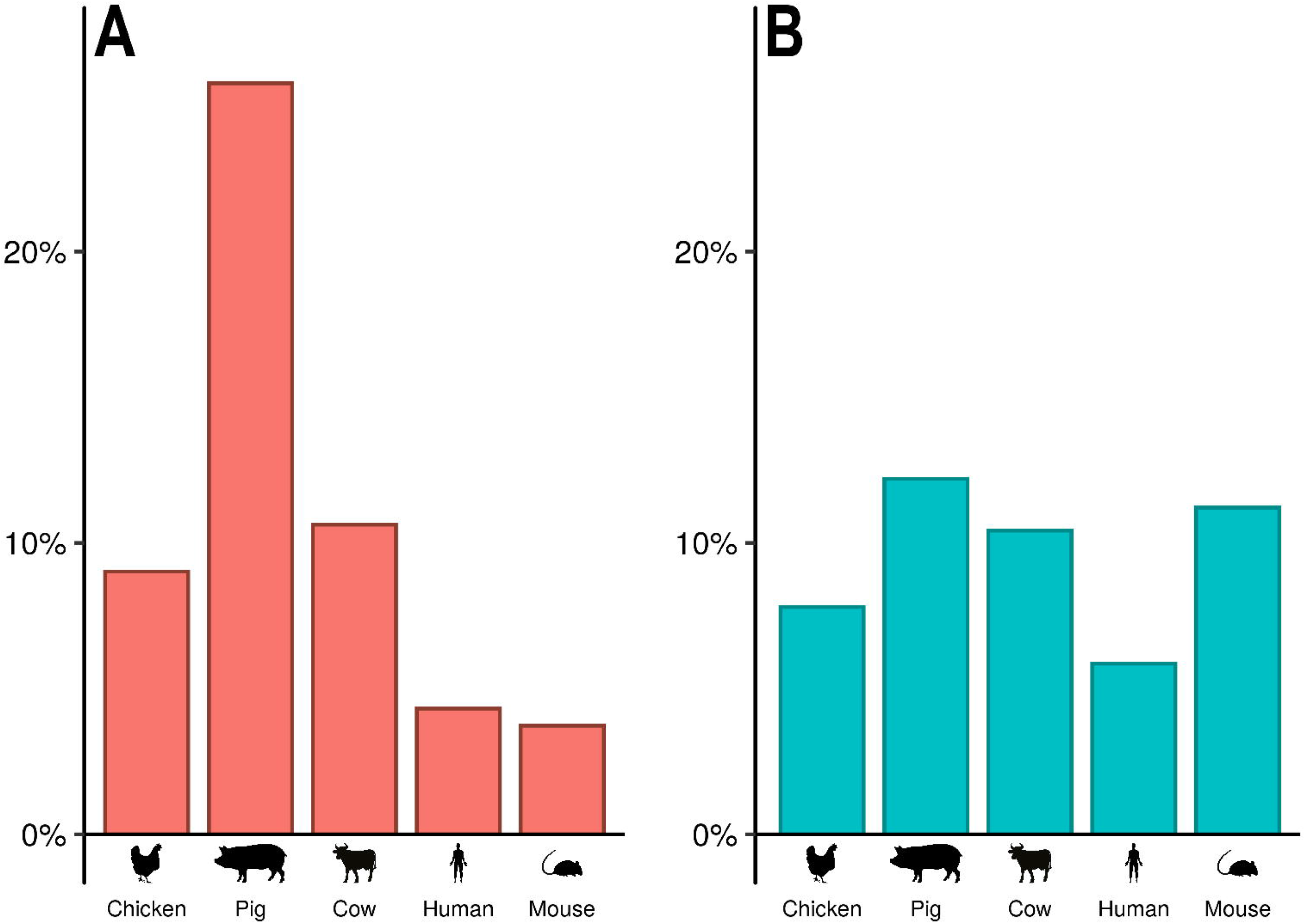
Proportion of re-assigned genomic position (panel A) and annotation (panel B) at probe-set level exemplified for chicken, pig, cattle, human and mouse. Results emphasize the increased accessible genomic knowledge which can be implemented in the revised assignments, *i.e.* chicken, pig, cattle, human and mouse microarrays.

The applied workflow combined genomic information and gene annotations which have been retrieved from various databases. Ranks can be used to define a specific priority for each database. Accordingly, the user can define the most actual and reliable database(s) as primary source for assignment and annotation. Nevertheless, the workflow allows to further implement other databases as well as own sequence information if available in a user-defined ranking scheme. This approach has proven to be beneficial in terms of the detectability of genes and transcript variants and the overall reliability of expression analyses [17].

## Implementation

The revised microarray workflow was implemented in the R package rePROBE (https://github.com/friederhadlich/rePROBE). rePROBE runs in the R environment under both Windows and Linux operation systems. It provides an easy-to-use text-based user interface. The *prepare_data* function comprises different R functions to automatically provide and prepare the needed information based on the initial array definition files. The interactive user interface queries basic requirements, including working directory, project name, species and the target databases. Subsequently, the revised probe-set assignment is prepared via the *run* function, which provides an R environment and data tables for downstream microarray analysis. A revised probe assignment and probe-set annotation for the ChiGene-1_0-st (*Gallus gallus*), PorGene-1_1-st (*Sus scrofa*) and BovGene-1_0-st (*Bos taurus*) microarrays are provided. The required computing time per species (15 CPUs, desktop-PC) is approx. 2-3h and includes indexing, probe assignment and probe-set annotation. A summary concerning the applied microarray platform can be accessed via the *show_report* function.

## Applications in transcriptional research

The application of the workflow generates information on genomic and exonic positions, polymorphisms, and probe assignment. In order to enable an interrelation with the vast majority of databases and genomic resources, the annotation refers to various transcript identifiers, including HUGO gene symbols and Ensembl IDs.

Regarding the outlined examples, the revised probe assignment and updated probe-set annotation has proven applicable for the ChiGene-1_0-st, PorGene-1_1-st and BovGene-1_0-st. In addition, the workflow for the HuGene-1_0-st and HT_MG-430_PM microarrays was successfully applied (see Supplemental table S1 for details). The workflow will contribute to describing a sophisticated picture of quantitative and functional traits in the species of interest. The physical redesign of microarray platforms will likewise improve the association between expression levels and corresponding transcripts [18]. Using microarray-derived transcriptional data as phenotypes will be beneficial in identifying expression quantitative trait loci (eQTLs) towards a sophisticated selection of candidate genes related to a trait in question [19,20].

The revised assignment according to up-to-date information will improve data quality as previously shown for different species [21]. However, the annotation level clearly depends on the probe-set design [22]. Probes are preferentially designed against the 3’ untranslated regions (UTRs) of transcripts. Respective sequence information is mostly derived by experimental evidence which does not necessarily represent the complete transcript sequence and may not be fully implemented in the current databases [22]. Therefore, the exclusion of unspecific probes according to the current database knowledge is indispensable as it substantially reduces the error due to cross-hybridizations at the probe level [23]. However, the partial mapping of probes especially represented by the number of genome-specific probes requires a sequential update of the assignment and annotation with a special emphasis on the growing knowledge of SNPs in the era of next-generation sequencing.

## Availability

rePROBE is freely available at GitHub (https://github.com/friederhadlich/rePROBE). All information regarding installation and application of the tool are provided.

## Author Contributions

FH, HR and MO conceived and designed the research. NT, EM, SP and KW reviewed the applied protocols. FH, HR and MO drafted the manuscript. NT, EM, SP and KW edited and revised the manuscript critically. All authors approved the final version of the manuscript.

## Acknowledgements

This work was partly funded by the Leibniz ScienceCampus Phosphorus Research Rostock and has received funding from the European Research Area Network on Sustainable Animal Production (ERA-NET SusAn) as part of the PEGaSus project (2817ERA02D). The Leibniz Institute for Farm Animal Biology (FBN) provided own matched funding.

## Supplementary material

Supplementary table S1 contains the summary of performance indicators referring to the applied exemplary microarray platforms, including chicken (ChiGene-1_0-st), pig (PorGene-1_1-st), cattle (BovGene-1_0-st), human (HuGene-1_0-st), and mouse (HT_MG-430_PM).

## References

[1] Suravajhala P, Kogelman LJ, Kadarmideen HN. Multi-omic data integration and analysis using systems genomics approaches: methods and applications in animal production, health and welfare. Genet Sel Evol 2016;48:38.

[2] Friggens NC, Blanc F, Berry DP, Puillet L. Deciphering animal robustness. A synthesis to facilitate its use in livestock breeding and management. Animal 2017;11:2237–51.

[3] te Pas MFW, Lebret B, Oksbjerg N. Invited review: Measurable biomarkers linked to meat quality from different pig production systems. Arch Anim Breed 2017;60:271–83.

[4] Edgar R, Domrachev M, Lash AE. Gene Expression Omnibus: NCBI gene expression and hybridization array data repository. Nucleic Acids Res 2002;30:207–10.

[5] Liu H, Bebu I, Li X. Microarray probes and probe sets. Front Biosci (Elite Ed). 2010;2: 325–38.

[6] Wernersson R, Nielsen HB. OligoWiz 2.0 - integrating sequence feature annotation into the design of microarray probes. Nucleic Acids Res 2005;33:W611–W615.

[7] Naraballobh W, Chomdej S, Murani E, Wimmers K, Ponsuksili S. Annotation and in silico localization of the Affymetrix GeneChip porcine genome array. Arch Anim Breed 2010;53:230–8.

[8] Milchevskaya V, Tödt G, Gibson T. A tool to build up-to-date gene annotations for Affymetrix microarrays. Genom Comput Biol 2017;3:e38.

[9] Sandberg R, Larsson O. Improved precision and accuracy for microarrays using updated probe set definitions. BMC Bioinform 2007;8:48.

[10] Wang Z, Gerstein M, Snyder M. RNA-Seq: a revolutionary tool for transcriptomics. Nat Rev Genet 2009;10:57–63.

[11] Veneziano D, Di Bella S, Nigita G, Laganà A, Ferro A, Croce CM. Noncoding RNA: current deep sequencing data analysis approaches and challenges. Hum Mutat 2016;37:1283–98.

[12] Müller F, Scherer M, Assenov Y, Lutsik P, Walter J, Lengauer T, Bock C. RnBeads 2.0: comprehensive analysis of DNA methylation data. Genome Biol 2019;20:55.

[13] Amaral ML, Erikson GA, Shokhirev MN. BART: bioinformatics array research tool. BMC Bioinform 2018;19:296.

[14] Kogenaru S, Yan Q, Guo Y, Wang N. RNA-seq and microarray complement each other in transcriptome profiling. BMC Genomics 2012;13:629.

[15] R Development Core Team. R: A language and environment for statistical computing. Vienna: R Foundation for Statistical Computing; 2008.

[16] Langmead B, Trapnell C, Pop M, Salzberg SL. Bowtie: An ultrafast memory-efficient short read aligner. Genome Biol 2009;10:R25.

[17] Yin J, McLoughlin S, Jeffery I, Glaviano A, Kennedy B, Higgins D. Integrating multiple genome annotation databases improves the interpretation of microarray gene expression data. BMC Genomics 2010;11:50.

[18] Marczyk M, Jaksik R, Polanski A, Polanska J. Affymetrix chip definition files construction based on custom probe set annotation database. In: Semantic Methods for knowledge management and communication, series studies in computational intelligence, Berlin Heidelberg: Springer;2011,135–44.

[19] Ponsuksili S, Murani E, Trakooljul N, Schwerin M, Wimmers K. Discovery of candidate genes for muscle traits based on GWAS supported by eQTL-analysis. Int J Biol Sci 2014;10:327–37.

[20] Ponsuksili S, Zebunke M, Murani E, Trakooljul N, Krieter J, Puppe B, Schwerin M, Wimmers K. Integrated Genome-wide association and hypothalamus eQTL studies indicate a link between the circadian rhythm-related gene PER1 and coping behaviour. Sci Rep 2015;5:16264.

[21] Dai M, Wang P, Boyd AD, Kostov G, Athey B, Jones EG, Bunney WE, Myers RM, Speed TP, Akil H, Watson SJ, Meng F. Evolving gene/transcript definitions significantly alter the interpretation of GeneChip data. Nucleic Acids Res 2005;33:e175.

[22] Ballester B, Johnson N, Proctor G, Flicek P. Consistent annotation of gene expression arrays. BMC Genomics 2010;11:294.

[23] Horn F, Nützmann HW, Schroeckh V, Guthke R, Hummert C. Optimization of a microarray probe design focusing on the minimization of cross-hybridization. Proceedings of the International Conference on Bioinformatics and Computational Biology 2011;1:3–9.

